# Polysynaptic signal propagation in networked neural masses

**DOI:** 10.64898/2026.04.29.721638

**Authors:** Varun Madan Mohan, James A Roberts, Anagh Pathak, Anthony M Harris, Caio Seguin, Andrew Zalesky

## Abstract

The routing of information across the brain’s structural network is central to its wide range of functional capabilities. However, the mechanisms underlying information routing in complex brain networks, particularly between regions that do not share a direct anatomical connection, remain poorly understood. Neural mass models (NMMs), a computational modelling framework capable of capturing complex neural dynamics across scales, can potentially be used to study the dynamical and network bases of these vital polysynaptic routing processes. In this study, we investigate polysynaptic signalling in three widely used NMMs, obeying Ornstein-Ü hlenbeck, Stuart-Landau, and Jansen-Rit dynamics, by tracking the propagation of a discrete, focal, high-amplitude perturbation across the underlying network. We find that polysynaptic propagation emerges in all tested NMMs when configured within dynamical regimes that effectively enhance the persistence of perturbations. We also find distinct parameter domains that maximise signal propagation to directly connected regions and to those separated from the source by at least two hops. Finally, we benchmark in silico stimulus propagation in the brain network against an empirical dataset of direct electrical stimulation trials, to explore the relative capabilities of the NMMs in capturing signal propagation to connected versus unconnected regions. This analysis highlights the significance of dynamical repertoire in capturing stimulus propagation outcomes. Overall, this study provides insights into how dynamical and network features shape signal propagation over complex brain networks.

## Introduction

The brain relies on the flexible transmission of information between regions, to perform its vast range of functions. Communication between brain regions is understood to evolve over the structural connectome – an intricate network of white matter tracts that form anatomical connections between regions [1, 2]. The brain’s structural network has a distinctly non-uniform architecture and lacks “all-to-all” connectivity [3–5]. This implies that information transfer between anatomically unconnected regions needs to evolve via polysynaptic, or “multi-hop” routes, where information is successively relayed across mutual neighbours before reaching the target.

Several lines of evidence point towards polysynaptic communication in the brain. For instance, regions that lack any measurable structural connectivity between them have been found to be functionally coupled, suggesting the involvement of additional regions in mediating information transfer [5, 6]. Polysynaptic communication is also understood to underlie the non-local effects of neurostimulation. For example, stimulating the left dorsolateral prefrontal cortex using transcranial magnetic stimulation (TMS), as a therapy for refractory depression, modulates the activity of the subgenual cingulate cortex, with which it shows no measurable anatomical connectivity [7]. Direct electrical stimulation also elicits responses in regions not directly connected to the stimulus site [8].

We still have a limited understanding of what principles dictate the routing of information across the brain’s intricate structural network, although several theories have attempted to address this gap from graph theoretic and biophysical standpoints [2, 9, 10]. While underlying principles can ideally be discerned through direct measurements of signal propagation, such approaches become challenging at the whole-brain scale. Computational modelling frameworks can prove useful in these scenarios, acting as “brain simulators” where activity can be manipulated and the effects tracked across regions simultaneously. Neural mass models (NMMs) replicate complex whole-brain phenomena through a combination of dynamical and network features, making this class of models a potential platform for studying neural communication and understanding how phenomena like polysynaptic routing are realised in the brain.

A neural mass model (NMM) is a mathematical description of the activities of dense collections of a large number of neurons, termed neural “masses”, and the interactions between them [11, 12]. They have been extensively used to further our understanding of brain function on various fronts, including the emergence of functional relationships [13–19], the link between neural mechanisms and electrophysiological/neuroimaging observations [18, 20–27], and neurological disorders [28–30], among several others. NMMs are typically written down as systems of differential equations that dictate how the activities of the masses, which might represent mesoscale neuronal populations or entire brain regions, evolve over time. In systems comprising multiple neural masses, individual activities are further shaped by network-mediated interactions. These interactions, which represent the communication between masses, are responsible for the emergence of global activity patterns and functional relationships. The variation of model parameters or the interaction structure can alter the evolution of the system and can be leveraged to simulate, for instance, neuropathological activities [31, 32], the impact of neurostimulation [33, 34], or other scenarios of interest. Previously proposed NMMs collectively showcase a wide range of complex dynamical behaviours including noisy fluctuations [15, 35], oscillations [20, 23, 26], and multistability [21, 36, 37], which have been used to model characteristic features of neural dynamics. However, these models remain largely untested in terms of whether they accurately capture neural communication characteristics. This can be evaluated by introducing a controlled perturbation, or stimulus, and tracking its propagation over the network of neural masses through the responses it elicits over time [34, 38, 39].

In this study, we explore the dynamical bases of polysynaptic routing of information in a complex brain network. We employ an in silico neurostimulation framework, where a brief high-amplitude pulse is introduced to the dynamics of a neural mass, and its effect on the rest of the network is assessed within a short time window. We approach this problem systematically, by evaluating three classes of NMMs of increasing dynamical complexity, in two network configurations. For each model, configured by a set of model parameters that are systematically varied, we carry out three analyses designed to evaluate various aspects of neural communication. The first determines whether multi-hop/polysynaptic communication emerges in NMMs, by tracking stimulus propagation along a simple chain motif. The second analysis assesses the spread of stimuli in a complex whole-brain tractography-derived network, across brain regions separated by increasing hops/topological distances from the source. The third and final analysis assesses whether the signal propagation in the models reflects empirically observed signalling, by gauging the similarity of modelled stimulus-response probabilities to those observed in empirical neurostimulation trials. Together, these three computational experiments allow us to evaluate the capacity of popular descriptions of neural dynamics in capturing patterns of polysynaptic signal propagation.

## Results

Our goal in this study was to characterise signal propagation in networks of neural masses exhibiting complex dynamics. We explored three classes of NMMs with increasing dynamical and biophysical complexity, namely, Ornstein-Ü hlenbeck (OU) [35], Stuart-Landau (SL) [40], and Jansen-Rit (JR) [23] (Fig.1A).

**Figure 1:**
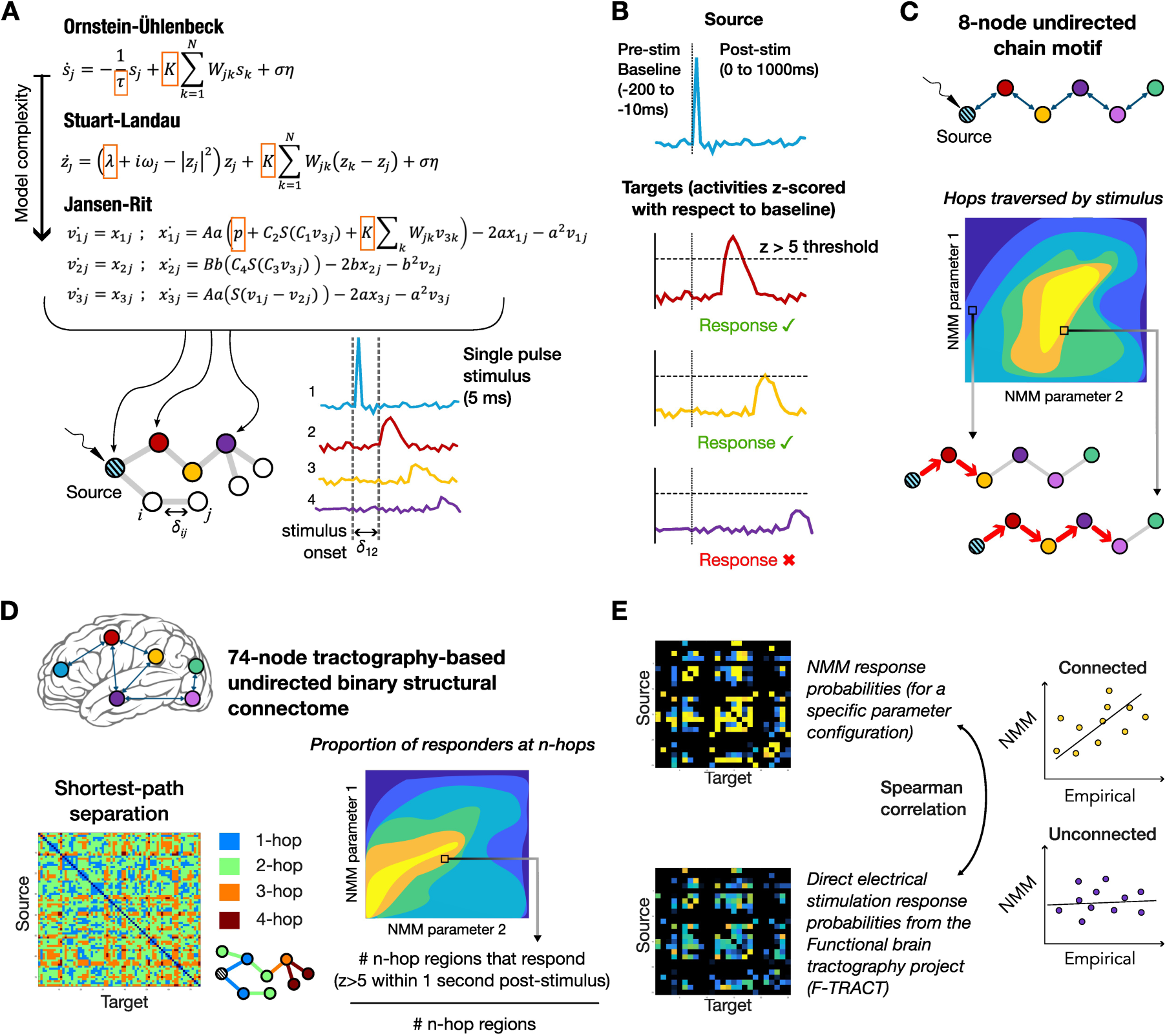
Analyses overview – **(A)** We analysed virtual stimulus propagation in networked systems of neural mass models (NMMs) exhibiting three classes of dynamics of increasing complexity. The parameters of the NMMs were identically set for each node in the network, and remained fixed, except for two parameters in each model, which were systematically varied (highlighted in orange boxes). We simulated a single-pulse stimulus in each node, for a duration of 5 ms, and tracked the changes in activities of nodes in the rest of the network. All networks were physically embedded, and a constant conduction velocity of 1 m/s was used to estimate signal transmission delays (*ω*) proportional to internodal separation **(B)** Stimulus propagation was tracked as per an empirical cortico-cortical evoked potential processing pipeline [42]. Briefly, activities were z-scored with respect to a [—200ms, —10ms] pre-stimulus baseline. If the z-scored activity in the post-stimulus window [0ms, 1000ms] crossed a threshold of |z| > 5, a response was recorded. **(C)** Our first analysis involved estimating the number of hops traversed by the stimulus in an 8-node undirected chain motif, with a single source at the start of the chain. 100 stimulation trials were carried out for each model configuration. **(D)** In the second analysis, the NMMs were embedded in a tractography-derived undirected binary structural connectome, and the proportion of responders at various separations from the source was assessed. **(E)** The response probabilities estimated from the NMMs, defined as the ratio of number of times a region responds (as per (B)) to the total number of stimulation trials, were compared to empirical estimates from the F-TRACT dataset. The correlations between directly connected (1-hop) and unconnected (2-to n-hop) region pairs were separately calculated. The connected and unconnected correlation coefficients across the tested parameter space were then rescaled and summed to identify the optimal model configuration that showed maximum empirical similarity in terms of both local and non-local responses to stimuli. 50 stimulation trials were carried out for brain network analyses.

For each of the dynamical classes, we initially assessed their ability to support multi-hop/polysynaptic signal propagation over a simple chain motif (Fig.1C), across a range of relevant model parameters. In our second set of analyses, the masses were embedded in a complex, tractography-based brain network (Fig.1D), allowing us to measure the length (in number of hops) of stimulus propagation paths in a biologically realistic network architecture. Finally, we sought to test whether the stimulus propagation characteristics in each dynamical model reflected the empirical propagation of direct electrical stimulation (Fig 1E). Using the same brain network setup as the previous analysis, we tested the correlation between measures of model-derived and empirical response to regional stimulation, the latter sourced from the Functional Brain Tractography (F-TRACT) dataset [41, 42]. Together, these three sets of analyses identify the parameter combinations and dynamical regimes in which polysynaptic signalling emerges in different NMMs.

### Stimulus propagation in a chain motif

Activities modelled by NMMs arise through a combination of two aspects: internal dynamics and network effects. To tease out these effects and assess the role of dynamical complexity (which varies across the tested NMM classes) in supporting polysynaptic propagation, we characterised how stimuli propagated over a bidirectionally-coupled chain motif, in the three models. The motif consisted of 8 neural masses that were equally spaced, exhibiting dynamics defined by a fixed parameter configuration. A single-pulse stimulus was then simulated at one end of the chain (the source), and the activities of all nodes were analysed to study its propagation across the connections. Specifically, we tested whether the activities of downstream nodes varied significantly from their respective pre-stimulus or baseline activities (Fig.1B). In this manner, we were able to estimate the average number of connections or hops stimuli traverse for a particular configuration of node dynamics. We systematically varied model parameters to delineate the parameter space where models exhibit multi-hop propagation down the chain motif. Importantly, the simple linear arrangement of the chain motif precludes contributions of converging and diverging network interactions, while offering a substrate for polysynaptic communication.

For OU dynamics (Fig.2A), we observed that increasing the global coupling, K, and the relaxation timescale (allowing for increased persistence of stimulus effects), *τ*, were both favourable for multi-hop propagation. The maximum number of hops observed was *≈* 3.37, indicating significant non-local signal propagation. This maximum was observed at the bounds of the tested parameter space, close to the stable limits of the dynamics (Suppl. Info., SI.S1).

**Figure 2:**
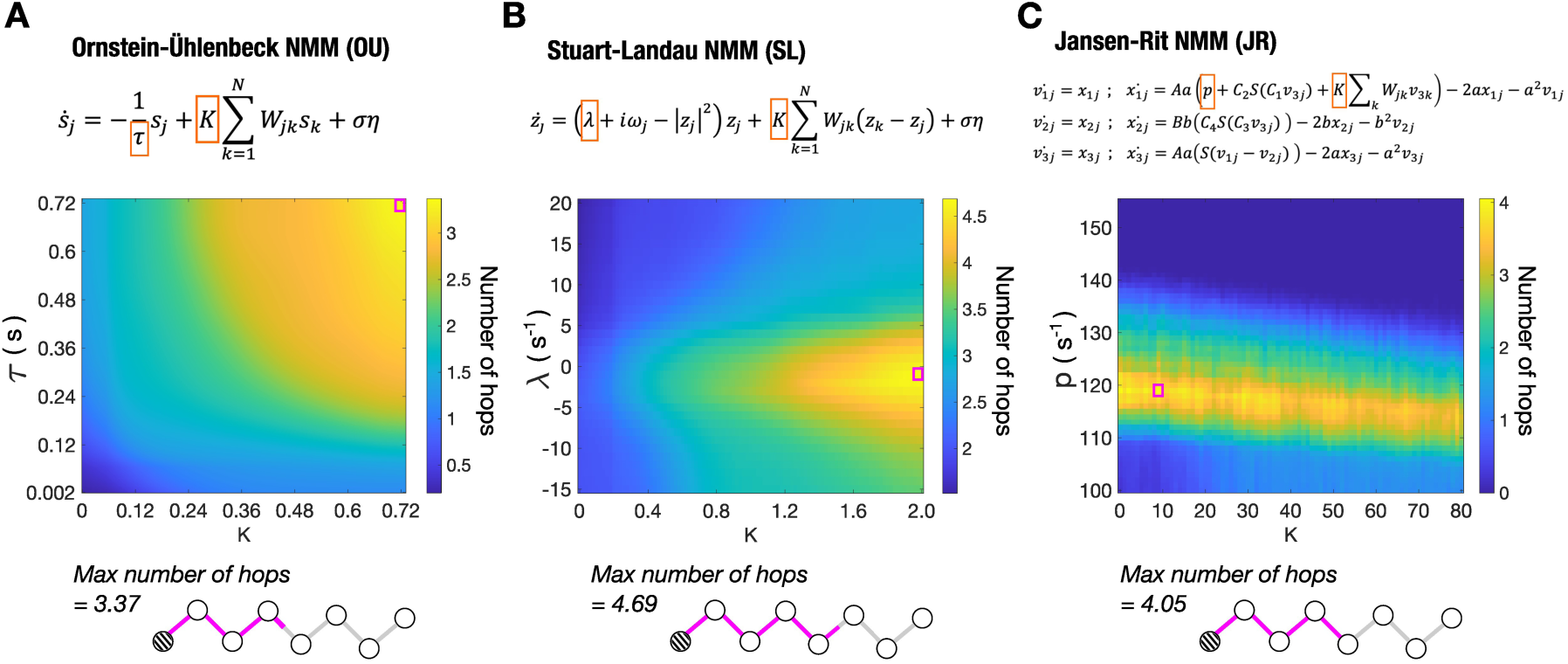
Multi-hop propagation in a chain motif. Varied model parameters are highlighted in orange in the differential equations governing the dynamics, in the top row. The parameter space maps present the average number of hops a stimulus propagated in an 8-node chain motif (100 stimulation trials) **(A)** Ornstein-Ü hlenbeck NMM. Varied model parameters: Timescale, *τ*, and the global coupling, K. **(B)** Stuart-Landau NMM. Varied model parameters: Bifurcation parameter, *λ*, and the global coupling. **(C)** Jansen-Rit NMM. Varied parameters: Average background firing rate, p, modelled as uniformly random noise input, and the global coupling. The maximum number of hops observed in the tested parameter space for each model (highlighted in the magenta box) is graphically depicted in the bottom row. The patterned node represents the stimulus source. All parameter space maps are smoothed with 2D moving averages for clarity.

In the case of SL (Fig.2B), an increase in multi-hop propagation was observed as *λ* approached 0—the critical value of the parameter delineating between limit-cycle and fixed-point dynamics—from both the positive and negative directions. This trend was observed as *λ* was varied for any given K. The average number of hops also increased with K, although the rate of change was dependent on *λ*, with the most pronounced change when *λ* = —1. Within the tested parameter space, SL exhibited clear multi-hop stimulus propagation, with a maximum average of *≈* 4.7 hops for *λ* = —1 at the maximum tested value of coupling, K = 2. We note that the maximum number of hops will increase beyond 4 and eventually cover the length of the chain as the coupling strength is increased beyond K = 2.

The JR models (Fig.2C), when configured with a background firing rate 110 s^—1^ < p < 135 s^—1^, displayed multi-hop propagation, across the tested K. For background firing rates outside this range, the maximum number of hops was close to 1, indicating a propagation to nodes only directly connected to the source, and showed little increase with global coupling. The maximum extent of propagation was *→* 4 hops for p = 119 s^—1^ and K = 10, indicating that stimulus propagation extended well beyond a source’s immediate neighbourhood.

In summary, across all models, we observed regimes of strong polysynaptic propagation over the chain motif as model configuration parameters were varied. Analysing the dynamical behaviour of the models indicated that the maximal extent of polysynaptic propagation was observed near bifurcation points (SI.S2). This suggests that model configurations that effectively increase the persistence of the brief stimulus are essential for the propagation of stimuli beyond the immediate neighbourhood of the source (see Discussion).

### Stimulus propagation in a brain network

In our second set of analyses, we considered stimulus propagation in a tractography-based brain network. The NMMs were initialised at each node of an unweighted binary structural connectome of a single hemisphere, in the Destrieux parcellation (N = 74 for a single hemisphere). The physical distances between nodes (gauged from the parcel centroids) were encoded into the NMMs as signal conduction delays. Prior to the analysis, we determined the shortest path separation between all region pairs from the structural connectivity matrix (Fig.1D). This revealed that the maximum separation in the network was 4-hops long. This step also helped determine the total number of 1-, 2-, 3-, and 4-hop separated pairs of regions. We then estimated the fraction of n-hop separated regions that responded to the stimulus. By systematically varying model parameters, we charted parameter spaces highlighting the dynamical regimes that may give rise to polysynaptic signalling chains in complex brain networks.

In OU, we observed that the share of responders 1-hop away from the source (regions directly connected to the source) increased as both the timescale and global coupling were increased, with a response elicited in over 90% of 1-hop regions when K > 0.08 and *τ* > 0.08 s (Fig.3A). A similar trend of increasing share of responders 2-hops from the source was observed as timescale and global coupling was increased, although the maximum share was only 65% (Fig.3B). Interestingly, we also noticed a reduced influence of K in increasing the share of responders when *τ* was low (< 0.06) i.e., when *τ* is low, increasing the coupling strength does little to increase the share of responders. This reduced influence of global coupling on the network involvement by stimuli was more apparent for the share of 3-hop separated responders, where we noticed a considerable effect of global coupling only at higher values of *τ* > 0.11 s (Fig.3C). The maximum share of 3-hop regions that responded was *≈* 40%, at the highest tested values of *τ* = 0.22 s and K = 0.22. Finally, we saw that the number of 4-hop responders increased as the timescale got longer but remained largely unchanged as global coupling was varied, apart from the variability stemming from the low number of 4-hop region pairs (Fig.3D).

**Figure 3:**
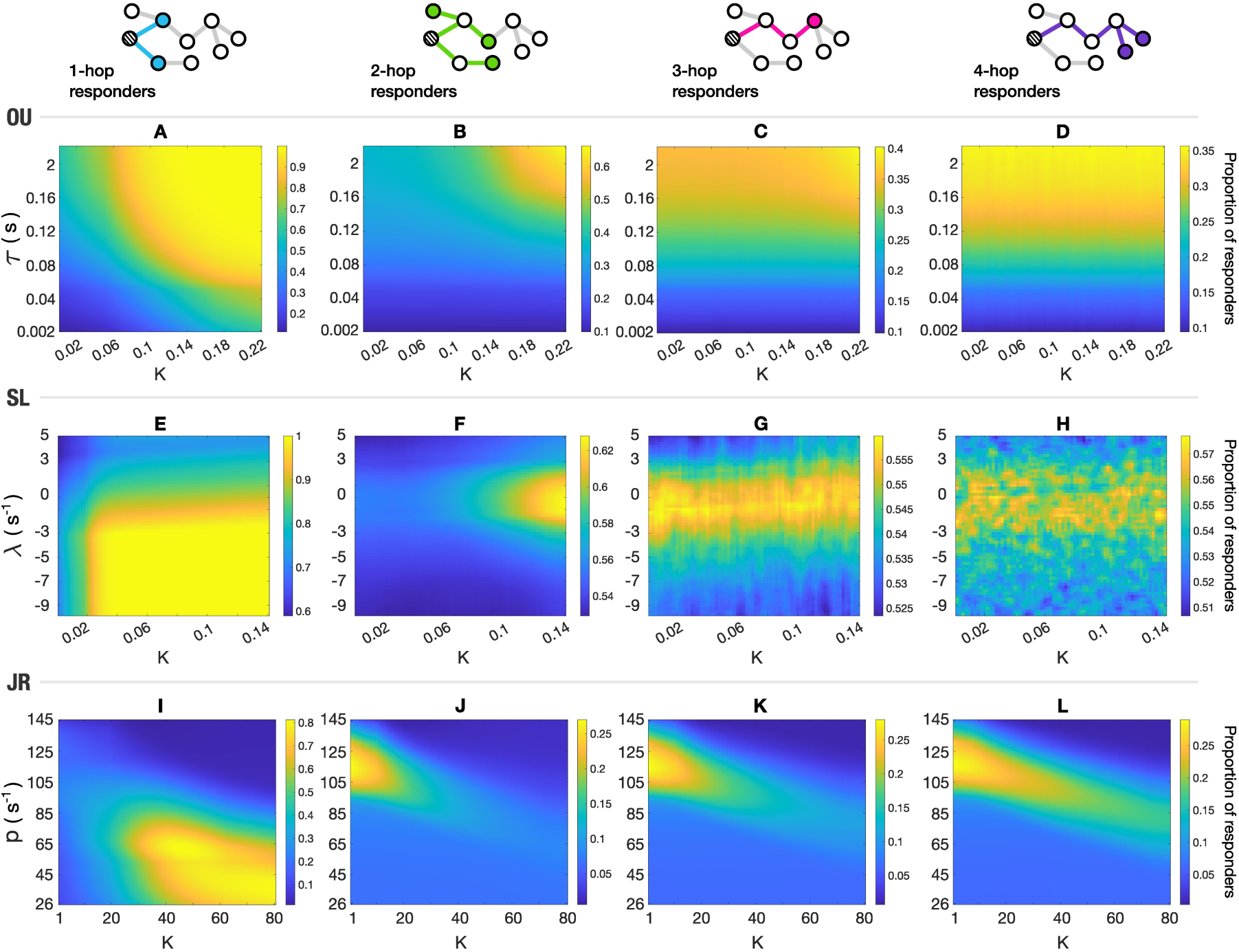
The proportion of responders (in a 74-node undirected whole-brain network) at increasing separations from the source (averaged over 50 stimulation trials). A schematic of n-hop neighbours is shown in the top row, with the patterned node depicting the source – the proportion of responders is estimated as the ratio of n-hop responders to the total number of n-hop neighbours. Note that the colour scales are set separately for each panel based on the range of observed values, to better showcase parameter-space variations; the maximum and minimum values that the responder proportion measure can assume are 0 (no responders) and 1 (all n-hop neighbours respond). Maps in each row correspond to each of the NMM classes (y-axis labels in A, E, and I apply to all panels in their respective rows). All maps are smoothed using 2D moving averages.

In the case of SL, specific parameter ranges maximised signal propagation to regions at increasing separations from the source. Specifically, in the tested parameter space, we found that 90%–100% of 1-hop or directly connected regions responded to the stimulus in the sub-critical to critical range, when *λ* < 0.5 and K > 0.02 (Fig.3E). We also found that over 50% of unconnected regions (2-to 4-hop) responded to the stimuli for all tested parameter configurations: *≈* 60% of 2-hop regions responded to the stimulus when *λ* was between —2 and 2 and K > 0.1 (Fig.3F), and *≈* 50 — 56% of 3-and 4-hop regions responded in models configured with *λ* between —3 and 1 (Fig.3G,H). Unlike OU, in which increasing both parameters resulted in increased shares of responders 1-to 4-hops away, SL configurations showing maximal shares of responders differed based on the separation considered, although polysynaptic propagation seemed to be maximal close to the critical value of *λ*.

For JR brain network nodes, we observed two distinct regimes supporting signal proportion to directly connected and unconnected regions. Configurations with background firing at the lower end of the tested firing rates, 35 s^—1^ < p < 65 s^—1^, and high global coupling, K > 40, showed a high percentage of connected regions responding to the stimulus (Fig.3I). However, less than 10% of unconnected regions responded for models within the same parameter range. The proportion of unconnected responders was higher at larger p ( > 85 s^—1^), with a maximum of *≈* 25% of 2 to 4-hop responders within the explored parameter space (Fig.3J,K,L). This was observed when p was close to 115 s^—1^—almost 2–3 times the background firing rate required for 1-hop responders.

To summarise, in this set of analyses, we embedded NMMs into a complex brain network and identified parameter regimes that supported stimulus propagation to connected (1-hop) and unconnected (2-to 4-hop) regions separately. These regimes showed limited overlap in general, implying that different dynamical behaviours optimise stimulus propagation to connected versus unconnected regions. Notably, polysynaptic propagation in the brain network was maximal around similar parameter regimes as in the chain-motif analysis, despite the marked difference in network scale and structure between the two.

### Alignment to empirical neurostimulation

In our final set of analyses, we benchmarked characteristics of virtual stimulus propagation in NMMs to those derived from empirical direct electrical stimulation in the F-TRACT dataset (Fig.1E). For each model configuration, the probability of a region responding to a stimulus was estimated for all source–target combinations. We then estimated the correlations between simulated and empirical response probabilities separately for regions with direct connections to the source (Fig.4A,D,G) and those that were 2 or more hops away (Fig.4B,E,H). Since a configuration that maximises empirical alignment for connected regions does not necessarily imply the same for unconnected regions and vice versa, the parameters that simultaneously maximised the correlation in both connected and unconnected regions was identified as the optimal model configuration. We hypothesised that stimulus propagation accuracy would increase with the dynamical repertoire of the NMMs, which would enable them to better capture the non-linearities shaping empirical neurostimulation outcomes.

**Figure 4:**
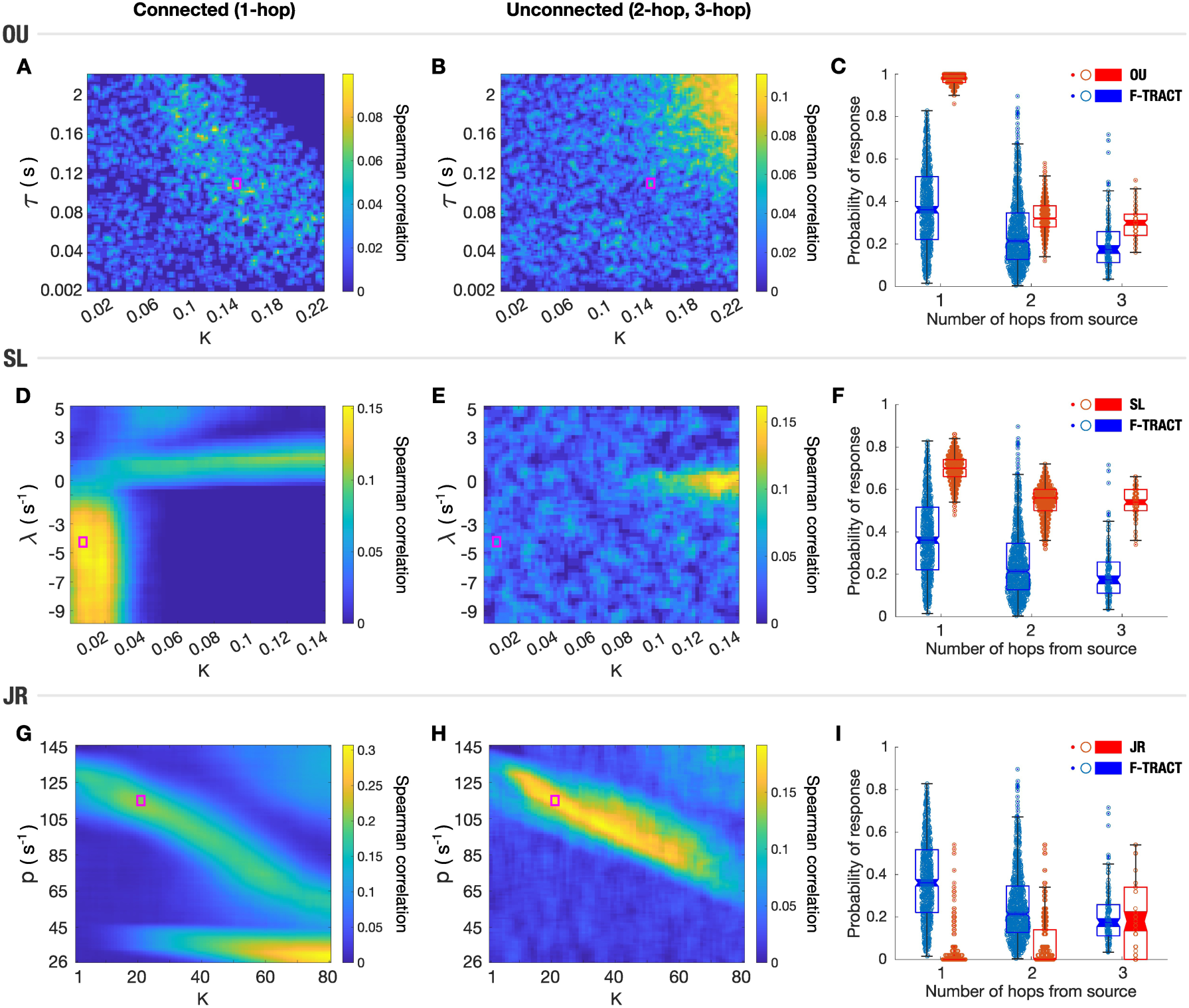
The correlation of NMM response probabilities to empirical estimates from the F-TRACT dataset. Plots in each row correspond to each of the NMM classes. **(A, D, G)** Correlation of response probabilities in connected (1-hop) regions. **(B, E, H)** Correlation of response probabilities in unconnected regions (2-and 3-hop). Only correlation values that passed a significance a significance threshold of 0.05 are reported. The optimal model configuration that maximised connected and unconnected correlations simultaneously is highlighted in the magenta boxes. **(C, F, I)** Distribution of empirical and simulated response probabilities. All maps are smoothed using 2D moving averages. The weak empirical correlation of response probabilities observed in OU across the parameter space is reflected in the noisy estimates even after smoothing.s

In our assessment of the OU model, we found that the response probabilities of directly connected regions were weakly correlated to those observed in F-TRACT across all model configurations, with a maximum Spearman correlation coefficient, R_sp_ *≈* 0.1 (p < 10^—4^) (Fig.4A). The correlation between the response probabilities of unconnected regions, specifically estimated to gauge the model’s polysynaptic signal transmission accuracy, showed an increasing trend with K and *τ*, and had a maximum of R_sp_ *≈* 0.11 (p < 10^—3^) (Fig.4B). The optimal model configuration that simultaneously maximised the correlation in connected and unconnected regions was for *τ* = 0.11 s and K = 0.14. Analysing the distribution of response probabilities with increasing hops from the source for this model, we found that regions that were directly connected to a source had a considerably higher probability to respond (median response probability *≈* 0.98) relative to empirical data (median response probability 0.36) (Fig.4C). Regions that were 2-or 3-hops away from the source were also more likely to respond on average, although this difference was much smaller compared to the directly connected regions (median response probability in NMM / F-TRACT; 2-hop: 0.32 / 0.21; 3-hop: 0.3 / 0.17). Moreover, the response probability distributions of 2-and 3-hop regions also showed more overlap with the empirical distributions.

Response probabilities in the SL model showed maximal correlation to F-TRACT within connected regions when configured in the subcritical non-oscillatory regime with *λ* < —2 and low coupling K < 0.03: The maximum observed correlation was R_sp_ = 0.152 (p < 10^—15^), for K = 0.018 and *λ* = —5.5 (Fig.4D). In contrast, models with *λ* < 0 and high coupling showed negligible empirical correlation, as a result of connected regions responding almost surely (response probability = 1) to stimuli. The empirical correlation of unconnected response probabilities was relatively high for NMMs with *λ ≈* 0 and K > 0.1 with a maximal correlation comparable to the connected regions – R_sp_ *≈* 0.162 (p < 10^—8^) at *λ* = —0.25 and K = 0.132 (Fig.4E). The model that maximised the correlation of both connected and unconnected response probabilities was configured with *λ* = —4.25 and K = 0.008. Comparing the response probability distribution for this model to empirical data, we found that regions at all separations were significantly more likely to respond in SL (Median response probability in SL / F-TRACT; 1-hop: 0.7 / 0.36; 2-hop: 0.56 / 0.21; 3-hop: 0.54 / 0.17) (Fig.4F). Relative to OU, however, 1-hop regions were less likely to respond, whereas 2-and 3-hop regions were more likely.

Finally, for the JR model, we found that the correlation between connected region pairs was stronger for low values of p and high K, with a maximum correlation of R_sp_ *≈* 0.3 (p < 10^—20^) observed for p = 30 and K = 80 (Fig.4G). In contrast, the correlation between unconnected regions was maximal for higher p and lower K, and the maximum correlation observed was R_sp_ *≈* 0.191 (p < 10^—6^), for p = 101 and K = 36 (Fig.4H). The model that simultaneously maximised the alignment in both connected and unconnected regions was found close to this regime, with p = 115 and K = 21. The probability-separation distribution for this model revealed that regions at 1 and 2 hop separations were significantly less likely to respond to the stimulus than in F-TRACT, whereas regions 3 hops away showcased median response probabilities that were empirically comparable (Median response probability in JR / F-TRACT; 1-hop: 0 / 0.36; 2-hop: 0 / 0.21; 3-hop: 0.18 / 0.17), unlike in OU and SL (Fig.4I).

Overall, across the models, we found that there were distinct dynamical regimes in which stimulus propagation was maximally aligned to empirical observations, depending on whether the target was directly connected to the source or not. As a result, the optimal model configuration, which considered the empirical alignment of responses in both connected and unconnected regions, did not coincide with the dynamical regime that only maximised polysynaptic communication (shown in the previous analysis). It was instead situated in an intermediate zone that supported propagation to both 1-hop and 2-to 4-hop regions.

In summary, our three-part analyses of the three NMMs revealed that each model exhibits multi-hop propagation only in specific parameter configurations. We also found that the configuration influences the relative number of responders at various separations from the source. Importantly, a configuration that maximises the number of responders at one separation might show minimal involvement of nodes at different separations, except in OU, where a high coupling and long timescale maximised participation of regions at all separations from the source. Benchmarking the responses of the NMMs to empirical observations revealed distinct parameter regimes that maximised model accuracy in terms of stimulus propagation to connected versus unconnected regions. In general, the NMMs were able to capture stimulus propagation to regions directly connected to the source better than to those that were 2-or 3-hops away, as signified by the differences in correlation to empirical signalling data. Finally, our analyses of the optimal models that simultaneously maximised the empirical correlation of connected and unconnected regions, revealed that JR showed the strongest correlation within the tested parameter range, followed by SL, and then OU (Connected / Unconnected R_sp_; JR: 0.205 / 0.186; SL: 0.149 / 0.063; OU: 0.087 / 0.037).

## Discussion

Understanding the dynamical processes by which quick, accurate, and efficient information routing is realised in the brain, particularly across polysynaptic paths, is an important problem in neuroscience. It has far-reaching implications for our understanding of structure–function relationships, dynamic communication, and the effects of neurostimulation [2, 43, 44]. In this work, we explored how polysynaptic signalling emerges in systems of interacting neural masses, and its relation to their underlying dynamics. We did this by studying how discrete, focal stimuli propagated in systems of in silico neural masses following Ornstein-Ü hlenbeck (OU), Stuart-Landau (SL), and Jansen-Rit (JR) dynamics. For each class of NMMs, we first evaluated their ability to showcase multi-hop (polysynaptic) propagation, and delineated model parameter configurations allowing the stimulus to engage nodes at various separations from the source. We then assessed the alignment of stimulus propagation in the NMMs to empirical neurostimulation outcomes and examined how the likelihood for a stimulus to elicit a response varied with the number of hops from the stimulation site.

Our initial analysis of polysynaptic propagation was carried out in a simple chain motif consisting of 8 nodes. Being the simplest architecture that supports polysynaptic propagation while minimising confounds of signal mixing through diverging and converging paths, the chain motif serves as an important reference to assess whether the dynamics fundamentally support signal propagation over multiple hops, in the absence of network effects. Our analysis revealed that polysynaptic propagation in the NMMs could only be realised for a limited range of parameters. This was for high values of coupling and long timescales for OU (Fig.2A), which intuitively sustains the perturbation and reduces its attenuation over hops. In the more complex SL (Fig.2B) and JR (Fig.2C), analysing their dynamical behaviour revealed that polysynaptic communication was maximal when the models were close to bifurcations; i.e., a regime where the behaviour of the system undergoes qualitative change—from non-oscillatory to oscillatory behaviour for SL, and from non-oscillatory to spiking behaviour for JR (SI.S2). It is likely that the propagation of the stimulus across multiple hops was facilitated by the increased sensitivity of regions to perturbations, which is characteristic of a system close to criticality [45, 46]. Additionally, it is possible that the phenomenon of “critical slowing down” [45, 47] effectively sustains the effects of the otherwise brief stimulus, allowing regions to respond more strongly, allowing for multi-hop propagation. Across the tested model configurations, we observed that the stimulus travelled the furthest in SL, *→* 4.7 hops, followed by JR, and finally OU.

In addition to obvious phenomena such as signal mixing, network effects can also effectively alter the dynamical behaviour. This is best demonstrated by the SL model. As per the differential equations governing SL dynamics, individual nodes, in the absence of any network effects, will transition to oscillatory behaviour when the bifurcation parameter, *λ*, exceeds 0. The chain-motif case indicates that multi-hop propagation is most apparent when the system is critical (*λ* = 0). The addition of network effects through the interaction term (modulated by the global coupling, K) however, effectively alters *λ*, proportional to the degree of the node; i.e., dynamics of nodes with a high degree will undergo bifurcation at a greater *λ* than those with lower degrees. Such effects add a connectivity-dependent complexity to NMMs, with potentially considerable implications to stimulus propagation (the change in *λ* affects the regime in which polysynaptic propagation occurs) (SI.S3). This further emphasises the importance of using simple network motifs for initial model assessment, to tease out dynamic-and network-specific contributions to stimulus propagation.

Our second set of analyses estimated the proportion of responding nodes at various separations from the stimulus source, in a system of 74 masses embedded in a human tractography-derived network. This captured the relative spread of the stimulus effects to directly connected (1-hop) and unconnected (2-to 4-hop) regions (Fig.3). Across models, we found that the parameter configurations maximising the involvement of nodes at different separations varied with the separation considered. In other words, configuring the model to show maximal involvement of 1-hop or directly connected regions, for example, did not guarantee a similar involvement of nodes n-hops away. By overlapping the responder proportion maps, we can identify the relative shares of responders at different separations from the source and effectively capture the multi-hop or polysynaptic profile of specific parameter configurations. As in the chain-motif case, we also note that stimuli elicit responses in unconnected regions the most when the models are tuned close to criticality.

Our final set of analyses involved correlating NMM response probabilities of connected (Fig.4A,D,G) and unconnected regions (Fig.4B,E,H) to empirical observations derived from the F-TRACT dataset. This primarily revealed that model configurations that maximise the similarity between responses of connected regions do not necessarily support empirically accurate response patterns in regions 2 and more hops away. Additionally, comparing the correlation maps to the responder proportion maps reveals that empirical correlation is generally lower for model configurations in which stimuli elicit responses in a large proportion of regions (> 90%). This is particularly evident in the OU and SL models, where we see an area of reduced correlation of connected response probabilities, overlapping the parameter domain associated with increased proportion of 1-hop responders. A reason for the reduced correlation is the high likelihood of responses, with minimal variability across region pairs. The F-TRACT-based empirical response probability distributions for regions separated from a source by 2-and 3-hops illustrates the prevalence of multi-hop or polysynaptic propagation in whole-brain communication. Although all tested NMM classes have parameter configurations supporting multi-hop stimulus propagation, the response probability distributions for the optimally configured models show a clear mismatch relative to the empirical distributions. The higher empirical correlation of JR relative to SL and OU, further suggests that out of the three models, JR might be better suited to applications that demand accuracy in terms of communication. It is, however, important to note that the magnitude of the empirical correlation was low across all models, indicating that despite being able to simulate empirically relevant multi-hop propagation, the tested NMMs fail to adequately capture other essential features of empirical stimulus propagation. Some possible reasons and directions to pursue to potentially improve the accuracy and predictive utility of the NMMs include:

1. The model parameters are identically set for all nodes; heterogeneity between nodes is only brought about by their respective locations in the physically-embedded network. Incorporating additional heterogeneity by setting each node’s fixed parameters, such as the intrinsic frequency, in an empirically-informed manner [34, 48] will likely enable models to better capture signal propagation characteristics.
2. Almost all conventional NMMs are formulated such that information (including stimuli) from a node is transmitted to all nodes it is directly connected to, scaled by the connection weight. Since the underlying connectivity network that defines the strength of interactions at any given time remains unchanged, regions continuously receive information from other networked masses. Introducing a biologically informed state-dependent interaction term would modulate the influence exerted on a mass by the rest of the network and potentially improve model accuracy with respect to stimulus propagation. A recent study reported significant improvements to model performance when dynamical interactions were implemented in a networked system of Kuramoto oscillators—the fixed amplitude variant of SL NMMs [49]. This approach would also be useful to assess theories of flexible communication [9, 50, 51] in silico, using NMMs. An alternate approach to realising diverse information transmission scenarios without invoking dynamic interactions is followed by network communication models, whose predictions were recently shown to yield strong correlations with the observations from F-TRACT [8].
3. All our assessments of model performance are based on response characteristics estimated through a cortico-cortical evoked potential (CCEP) processing pipeline originally used in the empirical F-TRACT data [42]. It is possible that certain features of the model that capture propagation might be masked by the CCEP pipeline, affecting response estimates. For instance, signal attenuation across multiple hops might be more pronounced in the NMMs than empirically, potentially leaving it undetected after CCEP processing. Adapting the CCEP pipeline to account for such model-specific features, such as a threshold proportional to a node’s distance from the source might be necessary to properly capture information flow in NMMs.

This work also serves to demonstrate the potential of NMMs as a platform to study neural communication mechanisms at the whole-brain scale. NMMs that feature biologically relevant signal propagation in addition to complex dynamics may also be applied as a “simulator” in domains such as neurostimulation, where systemic responses can be characterised in detail and effects of varying stimulation parameters and can be thoroughly analysed.

## Conclusion

In this work, we identify the dynamical regimes and parameter ranges in which NMMs give rise to polysynaptic signalling on top of connectome architecture. This will inform future work on modelling local routing mechanisms, and in discerning principles of inter-areal communication at the whole-brain scale.

## Methods

### Dataset

#### Structural Connectivity (SC)

The SC forms the anatomical substrate over which neural communication takes place and is central to modelling global dynamics using the NMM framework. In this study, we used a DWI-derived group-level SC (Destrieux atlas; 148 parcels) of 50 healthy young adults from the publicly available Microstructure Informed Connectomics (MICA-MICs) dataset [52], combined from subject-level SCs using the distance-dependent consensus algorithm [53]. The SC was proportionally pruned to retain the top 10% connections and then binarized. We only modelled the dynamics of a single hemisphere (N = 74), and so the SC was split, and the connectivity of the left hemisphere was used to constrain inter-regional interactions in the NMMs.

The signal transmission delays between each pair of regions (*ω*) were approximated by scaling the Euclidean distances between parcel centroids,D_ij_ as: *ω*_ij_ = D_ij_/v, where v is the conduction velocity, set to 1 m/s [54].

#### Functional Brain Tractography project (F-TRACT)

The F-TRACT dataset is a probabilistic atlas of cortico-cortical connections in humans, based on an extensive compilation of over 29000 focal direct electrical stimulation-response measurements from source-target pairs located across the cortical surface [41, 42]. The data is derived from intracranial electroencephalography (iEEG) measurements of the stimulation trials conducted in a cohort of 613 epilepsy patients (308 females), across multiple acquisition centres. Each stimulation run, which involves a pair of electrodes (source and target), typically comprised 30–40 stimulation pulses of 1 ms and 1–5 mA amplitude, with the actual parameters varying based on the acquisition centre, and was sampled at either 512 Hz or 1024 Hz. Electrode contacts were localised using pre-and post-implantation structural MRI and computerised tomography scans, and assigned to specific anatomical parcels according to various parcellation atlases; the Destrieux parcellation was used in this work [55]. Data from the F-TRACT project is publicly available in a group-averaged format at https://f-tract.eu/. We direct the reader to the publications that introduced the dataset [41, 42] for details on data acquisition, preprocessing, and quality control, but briefly cover processing steps relating to how responses were characterised.

#### CCEP processing

The responses over all pulses in a stimulation trial (satisfying quality criteria—see [42]) were initially averaged to yield the time course of the cortico-cortical evoked potential (CCEP). The CCEP was then z-scored with respect to a pre-stimulus baseline [—400, —10] ms. A stimulation was considered to have evoked a significant response if the z-scored CCEP crossed a threshold of z = 5 within 200 ms post stimulus. For each source and target parcel pair, the ratio of the number of trials with significant responses, to the total number of stimulation trials, defined the response probability. This yields an N *↑* N probability matrix, for a parcellation of N regions. To minimise inter-individual variability and measurement noise, we masked the response probability matrix such that only probabilities computed from a minimum of 200 stimulation trials were included.

The CCEP processing pipeline used in the F-TRACT dataset was used to derive the NMM response probabilities (variables capturing activities and responses are specified in respective NMM sections), except for the pre-stimulus baseline, which was set as [—200, —10] ms, in accordance with [42], and significant responses within 1000 ms following the stimulus were considered.

#### Neural mass models

Neural mass models are descriptions of the dynamics of neural masses—a population of a large number of neurons whose collective behaviour can be effectively described in terms of a small number of dynamical variables [11]. In this study, we explored signal propagation in NMMs defined using three classes of dynamics, namely, Ornstein-Uhlenbeck (OU), Stuart-Landau (SL), and Jansen-Rit (JR).

#### Ornstein-Ü hlenbeck

The OU process is one of the simplest descriptions of noise-driven, mean-reverting dynamics. Its behaviour is defined by two opposing components: a noise term that randomly fluctuates the system’s activity, and a restoring force term proportional to the system’s activity relative to a defined mean (zero in our case) which determines how quickly the activity reverts to the mean [35, 56]. Since the restoring force has a direct implication on how persistent the effects of the stimulus are, we fixed the noise amplitude for all our analyses, and systematically varied the inverse of the restoring force, *τ*, which intuitively captures the attenuation timescale of the OU process. For example, a low restoring force implies that a fluctuation from the mean, whether noise or stimulus driven, will take longer to revert. In addition to the timescale, we varied the global coupling parameter (K), which scales up the network’s connections, and increases the strength of interactions between the nodes.

The dynamics of the activity of a region j (at a time t) is defined as:

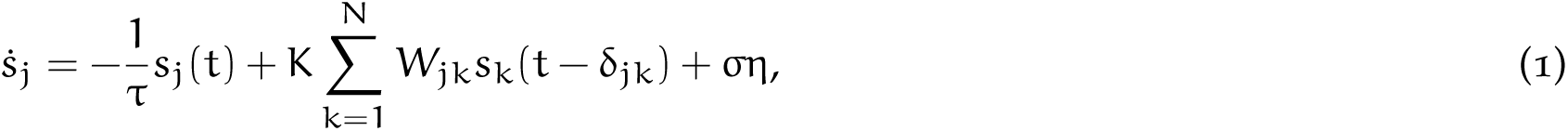

where W_jk_ is the strength of the connection between regions j and k, η is white noise, σ is the noise amplitude, and the *δ*_jk_ represents the signal transmission delay between the regions.

Stimuli were modelled as a discrete deviation in the activity, s, of a single region. Specifically, at the stimulus onset time (t = 0) the activity of the perturbation source was raised and maintained at 200 times the noise amplitude for a duration of 5 ms. Following the 5 ms perturbation, the activity of the source was allowed to evolve naturally for 1000 ms. CCEP processing was carried out on the time series of s.

#### Stuart-Landau

The second class of models we investigated was neural masses exhibiting Stuart-Landau dynamics (SL) with additive noise. The SL model describes variable amplitude phase oscillators [40] and adds a dimension of complexity over the OU model by allowing both phase and amplitude dynamics. Importantly, Stuart-Landau oscillators exhibit a bifurcation in dynamics, from noisy/asynchronous to oscillatory behaviour, upon variation of its bifurcation parameter, *λ*. Although considerably more complex than OU, the SL model is phenomenological and devoid of any explicit biophysical components. A region, j, following SL dynamics is described as a complex vector, z_j_, whose evolution is defined as:

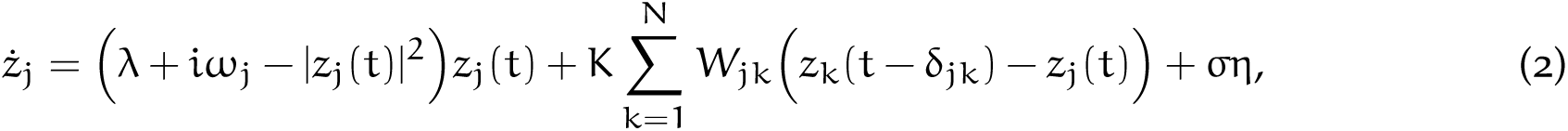

where *ω*_j_ is the intrinsic oscillation frequency of the region, |z_j_| is the amplitude of the oscillator, W_jk_ is the strength of the connection between regions j and k, η is white noise, σ is the noise amplitude, and the *δ*_jk_ represents the signal transmission delay. In all our analyses, the intrinsic frequency for all regions was fixed at 10 s^—1^, and the bifurcation parameter, *λ*, and global coupling strength, K, were varied.

The stimulus was modelled as a discrete and brief deviation of only the amplitude of the oscillator, |z|, whereas the phase was allowed to evolve naturally. This allowed the stimulus to be effective even in the sub-critical non-oscillatory regime of the SL model. As in the OU model, at the stimulus onset time (t = 0), the amplitude of the source region was raised and maintained at 200 times the noise amplitude for a duration of 5 ms, before being allowed to evolve naturally over the next 1000 ms.

The responses to stimuli were estimated by running the CCEP pipeline on the oscillator amplitudes, |z|, of the target regions.

#### Jansen-Rit

The Jansen-Rit (JR) model is a biophysically grounded model that describes the activity of a neural mass comprising interacting excitatory, inhibitory, and pyramidal subpopulations [23]. Its behaviour is defined by several parameters influencing timescales, couplings and gains of the subpopulations, and its functional complexity enables it to be configured to exhibit noisy, oscillatory and spontaneous spiking dynamics. The activity of a neural mass, j, is described by the second order dynamics of the average membrane potential v of each of the subpopulations, and written down as a set of differential equations of v and its derivative x for each of the subpopulations (1 *→* excitatory population, 2 *→* inhibitory population, 3 *→* pyramidal cell):

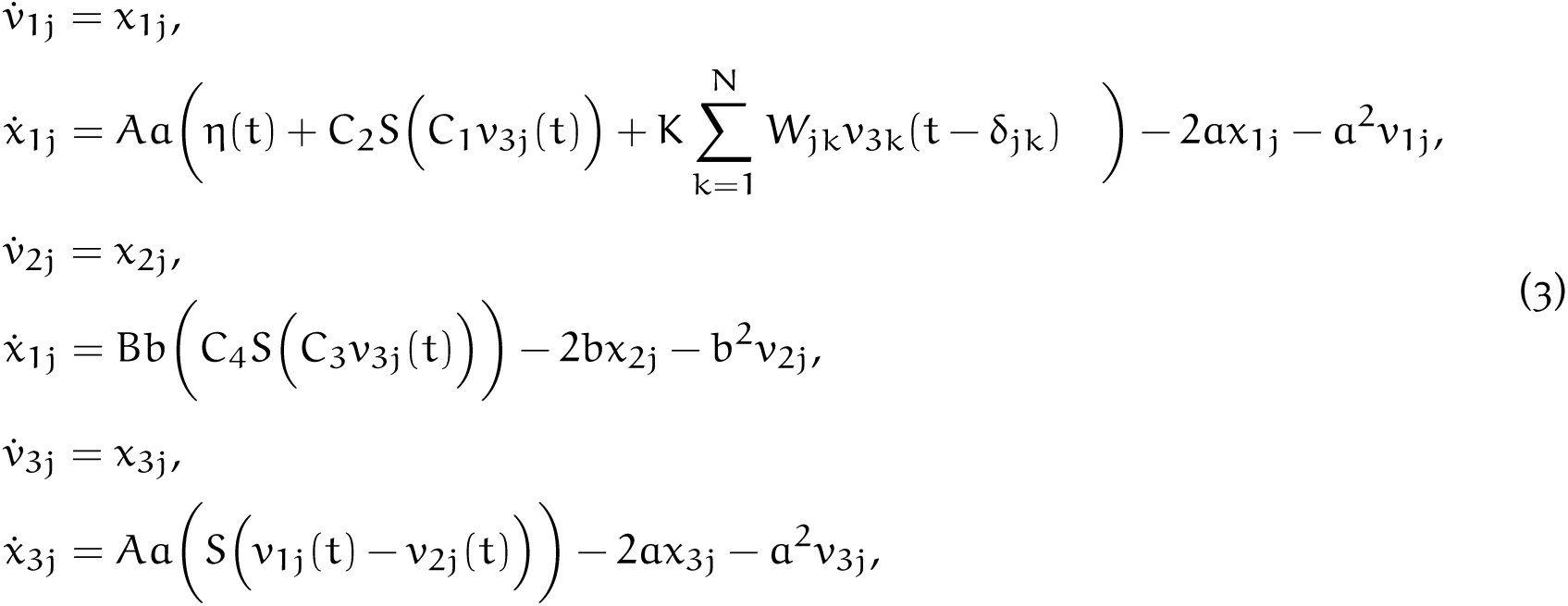

where *η*(t) is a random number representing the background firing rate to the excitatory subpopulation, sampled uniformly from an interval of width 50 s^—1^ centred around p. W_jk_ is the strength of the connection between regions j and k, and the *δ*_jk_ represents the signal transmission delay. S(v) is a nonlinear transfer function that transforms the average membrane potential of a population into an average firing rate, defined as S(v)= 2e_0_/[1 + e^r(v0—v)^], where e_0_ = 2.5 s^—1^, v_0_ = 6 mV, and r = 0.56 mV^—1^. The average number of synaptic contacts in the excitatory and inhibitory feedback loops are captured by C_1_ = C, C_2_ = 0.8C, C_3_ = 0.25C, and C_4_ = 0.25C, where C = 135. Other parameters include the average excitatory synaptic gain and the membrane time constants for the excitatory subpopulation: A = 3.25 mV and a = 100 s^—1^; and inhibitory subpopulation: B = 22 mV and b = 50 s^—1^. All these parameters were fixed for all regions, and across all simulations. For our analysis, we focused on the average background noise firing rate, p, which captures the spontaneous input into the system at baseline, and the global coupling, K. As a model with second order dynamics, The stimulation was modelled as an input to the excitatory subpopulation —specifically to ẋ _1j_ since the model has second-order dynamics—as a brief 5 ms long 1000 s^—1^ input added alongside p. As per convention, the output of a region was defined as the difference between the excitatory and inhibitory population potentials [23, 57]. The CCEP pipeline was applied to this variable to estimate responses.

#### Simulations

An important consideration for the extensive parameter space explorations carried out in this work was the computational optimisation of the simulation trials. Accordingly, all simulations were carried out in two phases: an initial stabilisation phase that allowed the dynamics to stabilise over 2 s, and a stimulation phase, in which each region was chosen as a stimulus source, and the dynamics of the network was allowed to evolve for 1 s after the stimulus duration of 5 ms. In SL, certain models configured in the supercritical, oscillatory parameter regime were found to take longer than 2 s to stabilise; models with these configurations were selectively allowed to stabilise over 20 s before initiating the stimulation phase. After each stimulation, the next source region is chosen, and the state variables are “reset” to the values at the end of the stabilisation phase, before running the stimulation trial with the new source. After all regions were considered as sources, the CCEP pipeline was initiated to assess the binary response matrix that identified whether the stimulus from a source was able to evoke a significant response in a target. Following this, another simulation was initiated with random initial conditions – each new simulation trial required a new run of the stabilisation phase before running the stimulations across regions. For each parameter configuration, we ran 50 simulation trials. All NMMs were numerically integrated using the modified Euler / Heun scheme, with a timestep of 10^—4^ s.

## Acknowledgements

V.M.M. was supported by the Melbourne Research Scholarship, University of Melbourne. A.P. acknowledges support from the Fondation Vaincre Alzheimer. A.M.H. was supported by the Australian Research Council (DE220101019). C.S. acknowledges support from the Australian Research Council (DP170101815). A.Z. is supported by an ARC Future Fellowship (FT220100091) and the Rebecca L. Cooper Foundation. This research was supported by The University of Melbourne’s Research Computing Services and the Petascale Campus Initiative.

