## Supplementary information for "Polysynaptic signal propagation in networked neural masses"

### S1. *Stability of Ornstein-Ühlenbeck dynamics*

The parameter ranges explored in the main text for the OU model have upper limits of $K = 0.22$ and $\tau = 0.22$. The stable parameter regime for the OU model is derived below.

The OU dynamics of a region $i$, embedded in a binary network $A$, with a global coupling $K$ and timescale $\tau$, is given as:

$$\frac{dx_i}{dt} = \frac{-1}{\tau}x_i + \sum_j W_{ij}x_j + \sigma\eta. \tag{1}$$

We can set the noise amplitude $\sigma = 0$, for simplicity. The above equation can be cast in vector form, as

$$\begin{aligned}\frac{d\mathbf{x}}{dt} &= \frac{-1}{\tau}I\mathbf{x} + KW\mathbf{x},\\ &= \left(KW - \frac{1}{\tau}I\right)\mathbf{x}.\end{aligned} \tag{2}$$

Integrating with a timestep of $h$,

$$\begin{aligned}\mathbf{x}_{t+1} &= \mathbf{x}_t + h\left(KW - \frac{1}{\tau}I\right)\mathbf{x}_t,\\ \mathbf{x}_{t+1} &= B\mathbf{x}_t,\end{aligned} \tag{3}$$

where $B = I + h\left(KW - \frac{1}{\tau}I\right)$. The stable parameter regime is defined when $max\left(\left|eig(B)\right|\right) < 1$.

For the tractography-based binary adjacency matrix used in this work, and $h = 10^{-4}$, we see that the stable regime is delineated below. The regime used in the main text is highlighted in the red box.

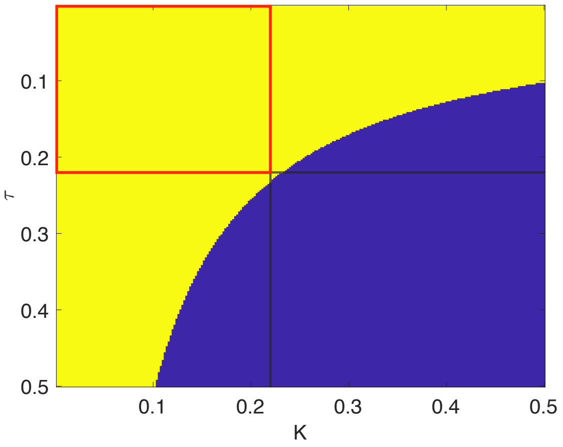

### S2. *Criticality in the Stuart-Landau and Jansen-Rit models*

*Jansen-Rit*

The figure shows the maximum and minimum value of the difference between excitatory and inhibitory subpopulation membrane potentials (after stabilisation of the dynamics), as a function of the average background noise firing rate. The global coupling, $K = 1$. For each $p$, 10 simulation trials were carried out. Models were randomly initialised at each trial. A transition from non-oscillatory to a spiking regime is observed at $p = 114$. Another transition is observed around $p = 140$, from spiking to a limit-cycle oscillatory regime.

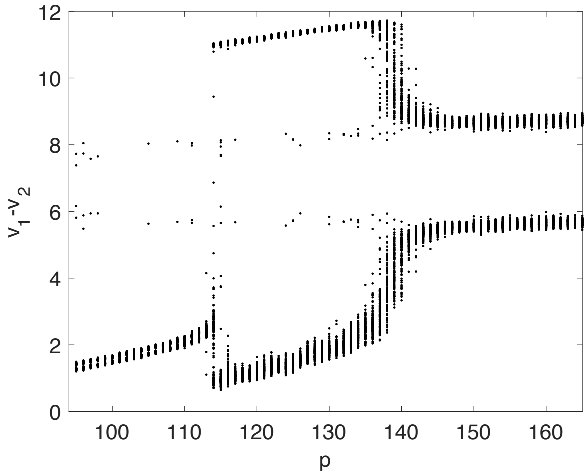


*Stuart-Landau*

The figure shows the amplitude of the Stuart-Landau oscillator as a function of the bifurcation parameter, $\lambda$. $K = 0.01$. For each $\lambda$, 10 simulation trials were carried out. The increase in amplitude after $\lambda = 0$ marks a bifurcation from non-oscillatory behaviour to a stable limit cycle with an amplitude of $\sqrt{\lambda}$.

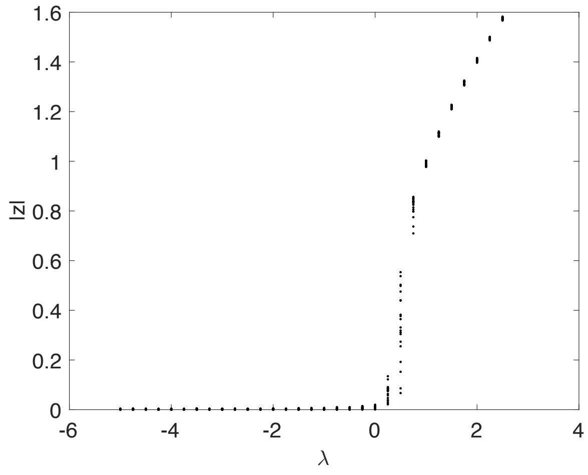

### S3. *Network effects on the Stuart-Landau bifurcation parameter*

The activity of a region j embedded in a network $W$ and oscillating with an intrinsic frequency $\omega$, following Stuart-Landau dynamics, is given by (we can disregard delays and noise for simplicity):

$$\dot{z}_j = \left(\lambda + i\omega_j - |z_j|^2\right) z_j + K \sum_{k=1}^{N} W_{jk}\left(z_k - z_j\right). \tag{4}$$

In polar form,

$$\dot{z}_j = \dot{r}_j e^{i\theta_j} + r_j i e^{i\theta_j} \dot{\theta}_j. \tag{5}$$

Expanding $z_j$ and separating $\dot{r}_j$ and $\dot{\theta}_j$, we obtain:

$$\begin{aligned}
\dot{r}_j &= (\lambda_j - r_j^2) r_j + K \sum_k^N W_{jk}(r_k \cos(\theta_k - \theta_j) - r_j), \\
&= (\lambda_j - r_j^2) r_j + r_j K \sum_k^N W_{jk}\left(\frac{r_k}{r_j} \cos \Delta\theta_{jk} - 1\right), \\
&= \left(\lambda_j - r_j^2 + K \sum_k^N W_{jk}\left(\frac{r_k}{r_j} \cos \Delta\theta_{jk} - 1\right)\right) r_j, \\
\dot{r}_j &= (\lambda_j' - r_j^2) r_j,
\end{aligned} \tag{6}$$

where $\lambda_j'$ is the effective bifurcation parameter of the region, given the network structure and inputs:

$$\begin{aligned}
\lambda_j' &= \lambda_j + K \sum_k^N W_{jk}\left(\frac{r_k}{r_j} \cos \Delta\theta_{jk} - 1\right), \\
&= \lambda_j + K \sum_k^N W_{jk}\left(\frac{r_k}{r_j} \cos \Delta\theta_{jk}\right) - K \sum_k^N W_{jk}.
\end{aligned} \tag{7}$$

where the last term, $K \sum_k^N W_{jk}$, is the degree (or strength) of node j.
